# Organic compounds drive growth in phytoplankton taxa from different functional groups

**DOI:** 10.1101/2023.10.06.561152

**Authors:** Nele Martens, Emilia Ehlert, Widhi Putri, Martje Sibbertsen, C.-Elisa Schaum

## Abstract

Phytoplankton are usually considered autotrophs by default, but an increasing number of studies shows that many taxa are able to also utilise organic carbon. Acquiring nutrients and energy from different sources might enable an efficient uptake of required substances and provide a strategy to deal with a varying resource availability, especially in highly dynamic ecosystems such as estuaries. In our study we investigated the effects of 31 organic carbon sources on the growth of 17 phytoplankton strains from the Elbe estuary spanning four functional groups. All of our strains were able to make use of at least 1 and up to 26 organic compounds for growth. Pico-sized green algae such as *Mychonastes, Choricystis* and *Chlorella*, as well as nano-sized green algae from the genus *Monoraphidium* in particular were positively affected by a high variety of substances. Reduced light availability, typically appearing in turbid estuaries and similar habitats, resulted in an overall poorer ability to utilise organic substances for growth, indicating that organic carbon acquisition was not primarily a specific strategy to deal with darkness. Our results give further evidence for mixotrophy being an ubiquitous ability of phytoplankton and highlight the importance to consider this trophic strategy in research.

## Introduction

Phytoplankton may play a more complex role in the carbon cycle than previously assumed. While one of their main roles is fixing CO_2_ through photosynthesis -which is the basis of carbon sequestration (1) -various studies now provide evidence that many taxa are actually able to acquire organic carbon from their environment, either by phagotrophy (2,3) or by uptake of dissolved compounds (4,5). Mixotrophy appears for taxa from miscellaneous functional groups of phytoplankton, including haptophytes (2,4,6), green algae (5,7,8), dinoflagellates (9), cyanobacteria (10), cryptophytes (11) and diatoms (12).

Combining autotrophic and heterotrophic mechanisms might enable an efficient uptake of carbon and required nutrients such as N, P or amino acids (13,14). Mixotrophic phytoplankton can achieve higher biomass yields in the presence of organic carbon compared to autotrophic conditions (15,16), which is made use of in bioengineering (17). The availability of different pathways of energy and nutrient acquisition might also have substantial benefits in highly dynamic ecosystems, e.g. when light, prey or nutrient availability vary (4,6,9,18), and allow phytoplankton to be a stable food source for zooplankton (19).

In tidal estuaries water masses are constantly reshuffled while phytoplankton drift between freshwater and saltwater ecosystems. Here phytoplankton are exposed to high variations in environmental conditions such as salinity and turbidity (20), as well a high number of organic substances such as amino acids (21) or fatty acids (22). The utilisation of organic compounds in such ecosystems might be critical for phytoplankton to maintain growth, especially where light is limited as a consequence of high loads of suspended matter. Different phytoplankton taxa have been shown to survive and grow based on organic carbon in the dark and at reduced light levels (4,9,23–25). Hence, unsurprisingly, the importance of mixotrophic taxa in estuarine ecosystems has been reported in several studies (26–29).

There is an increasing awareness that mixotrophy of phytoplankton might be rather the norm than the exception (13,30). This alters our understanding of food webs and substrate cycles. As part of the microbial loop as well as by grazing on bacteria and other small organisms, mixotrophic phytoplankton contribute to trophic upgrading (31). Here, they do not only pass on energy and nutrients as such, but might also alter them to become more beneficial for zooplankton (32), e.g. by providing high lipid and protein contents (17). By incorporating both organic and inorganic matter, biomass of mixotrophic phytoplankton becomes decoupled from the actual primary production. Mixotrophic phytoplankton can have a reduced chlorophyll content (7,15,33) and might therefore be underrepresented quantitatively with conventional measuring techniques that make use of pigment concentrations. Ultimately, recent studies show that integrating mixotrophy in climate change research is crucial for understanding carbon dynamics (34,35).

Here, we investigated the effects of 31 dissolved organic compounds on the growth of 17 phytoplankton strains isolated from the Elbe estuary by using Biolog EcoPlates™. Our aim was to compare the potential of different phytoplankton taxa from different functional groups to utilise dissolved organic carbon for growth. We moreover wanted to investigate the effects of reduced light availability that might occur frequently in the turbid estuary.

## 1 Methods

### Isolation and cultivation of phytoplankton

Water samples for the isolation of phytoplankton were collected in March (2021-03-12), May (2021-05-07/08) and February (2022-02-28) on the research vessel Ludwig Prandtl (LP210308, LP210503, LP220228) as well as during one sampling from the pier in July (2021-07-21). The sampling stations were located at 610 km, 634 km and 690 km distance from the spring of the Elbe river in Czech Republic. Additional information about abiotic parameters during sampling is given in the supplementary data (tab. S1).

We used a dilution approach to isolate the phytoplankton strains. This was conducted either in 96 well plates, where samples were diluted to contain on average 0.5 cells/ well or on agarose by picking colonies grown from single cells. Each strain went through this process at least twice to ensure clonality. We obtained 17 clonal phytoplankton strains (fig. 2, tab. S1). Note that isolation success is biassed by the abundance of taxa but also by their viability in the laboratory and does not necessarily reflect the natural communities. We did not remove the microbiome, as former studies have shown that phytoplankton can depend on bacteria in their environment (8,36). Moreover, co-occurrence and interactions with bacteria reflect the natural conditions in the field.

All strains were maintained in WHM freshwater media to which silicate was added (3 mg Si/L). The pH of the media was approximately 7. In a common garden approach, winter strains were kept at 12°C respectively 15 °C and summer strains at 18 °C. Strains were held at a 12:12 light:dark cycle at approximately 150 μEinstein/(s m2) in the light phase and gently mixed at 60 rpm.

All included strains are shown in fig. 2 as well as in the supplementary data (tab. S1) together with further information (e.g. origin, abiotics).

### Identification of the strains

DNA was extracted using a CTAB protocol (37). Cyanobacteria were identified by 16S sequencing (27F forward 5’-AGAGTTTGATCCTGGCTCAG-3’, 1429R reverse 5’-GGTTACCTTGTTACGACTT-3’) and eukaryotes were identified by 18S sequencing (5’-GCTTGTCTCAAAGATTAAGCC-3’ forward, 5’-GCCTGCTGCCTTCCTTGGA-3’ reverse) using the ncbi BLAST database. Additionally, morphological criteria from microscopy (Keyence BZ-X800) and flow cytometry (BD accuri C6 plus) were included to characterise the different strains. Green algae were assigned to the pico green or nano green fraction based on their size represented by their flow cytometric characteristics (see supplementary data tab. S1).

Our strains were identified as seven pico green algae, six nano green algae, three cyanobacteria and one diatom (tab. S1). All strains differed from each other either morphologically, with respect to their origin and sampling season and/ or genetically, hence representing unique geno- or ecotypes. Further information about how we assigned the taxa to the different strains is provided in the supplementary data (tab. S1).

### Experimental setup

An overview of the experimental setup is given in fig. 1a. We used Biolog EcoPlates™ to analyse the effects of organic carbon sources on the growth of the phytoplankton strains. The plates contained 31 organic substances as well as a control without an organic compound in triplicates. Organic compounds included 10 carbohydrates, nine carboxylic acids, six amino acids, four polymers and two amines. We added 100 μL of culture to each well of the EcoPlates™.

**Fig. 1:**
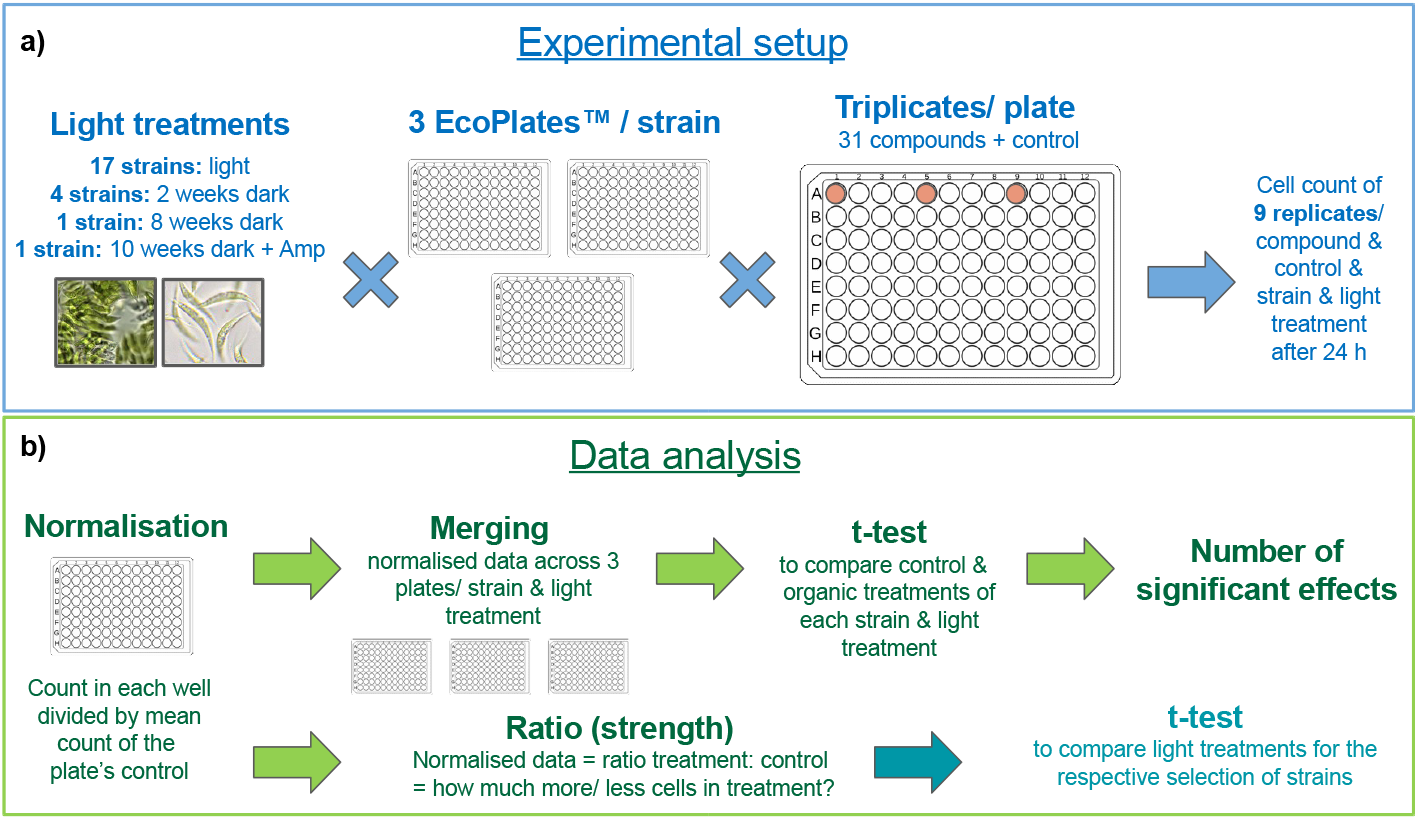
Experimental setup (a) and data analysis (b). ‘Light’ and ‘dark’ refer to the condition of the light phase of the 12:12 light:dark cycle, which were ca. 150 μEinstein/(s m2) in the light treatment and ca. 75% reduced in the dark treatment. Amp = Ampicillin (100 μg/ mL) added ca. five days prior to measurements

After approximately 24 hours, we used flow cytometry (BD accuri C6 plus) to determine the number of cells in each well. Here, we measured 10 μL per well with a flow rate of 66 μl/ min, regular cleaning and mixing between the samples. Gating was conducted optically based on the obvious phytoplankton clusters. The cell count after ca. 24 h incubation in the plates was used as a proxy for growth. While this is not a perfect proxy for growth, we refer to it as ‘growth’ throughout the text for better readability.

All 17 strains were included with standard light incubation, i.e. 12:12 light:dark cycle, where the photon flux in the light phase was ca. 150 μEinstein/(s m2). We refer to this setup as ‘light’ throughout the study. In the strains habitat, light availability quickly declines along the water column due to high turbidity (38). We therefore wanted to also investigate the effects of reduced light availability on the use of organic carbon for growth. For practical reasons, we selected a subset of four strains that particularly covered the largest groups of our pool of strains -i.e. nano and pico greens. Those strains included one strain of *Mychonastes* (strain P4), *Choricystis, Tetradesmus* as well as one strain of *Monoraphidium* (strain N7). Light reduction across wavelengths was achieved by semi-transparent foil (Lee Zirkon Filter, type 210), while running the standard 12:12 light:dark conditions of the incubators as described above. We refer to the ca. 75 % light reduction as ‘dark’ or ‘darkness’ throughout the text. The strains were kept in the dark for two weeks prior to the measurements, then transferred to the EcoPlates™ and incubated in the dark in the presence of the organic compounds. To test for the effects of bacteria and longer exposure to darkness, we additionally ran the experiment with *Mychonastes* (strain P4) with a dark acclimation time of eight weeks as well as by adding Ampicillin. The latter was done after ten weeks in darkness and Ampicillin was added as 100 μg/ mL ca. five days prior to the measurement. During the longer phases in the dark, i.e. more than two weeks, cultures were regularly transferred to fresh media.

To cover potential plate effects, we repeated the experiment three times for every strain and light treatment, including different densities within the exponential growth phase. This resulted in a total of ca. 70 plates included in this study. Exponential growth was ensured by casual tracking of the cell counts prior to the transfer of the cultures to the EcoPlates™ (see supplementary data fig. S1).

Metadata for the measurements -e.g. exact incubation times -can be found in the supplementary data (tab. S2).

### Statistical analysis

Cytometric data (cell counts) were processed and analysed in R (version 4.1.3), including the packages tidyverse (version 1.3.2) and stats (version 4.1.3). We used ggplot2 (version 3.4.0) for graphical presentation of data and LibreOffice Draw (version 7.1.2.2) for overview figures and addition of text notes.

An overview of the data processing is shown in fig. 1b. Results of the plate replicates were similar despite different cell densities, allowing to merge data across plates for every strain and light treatment. To do so, we first normalised the data of each plate by the plate’s control mean and then merged the datasets, resulting in nine replicates per strain, light treatment and organic compound respectively control. We then used the t-test (R, package stats version 4.1.3) to assess whether there were significant differences between each treatment with different organic compounds and the control (p ≤ 0.05, see supplementary data tab. S3). This gave us the number of significant positive and negative effects per strain and light condition across organic compounds, i.e. how many compounds positively or negatively affected our strains (fig. 2a+b). The normalised cell counts i.e. ratios of organic treatments vs. controls were moreover used to describe the strength of the effects on the strains, i.e. how much higher or lower cell counts were compared to the control (fig. 2c). For clarity we subtracted 1 from all ratios so that negative effects are represented by negative values and positive effects by positive values (fig. 2c).

**Fig. 2:**
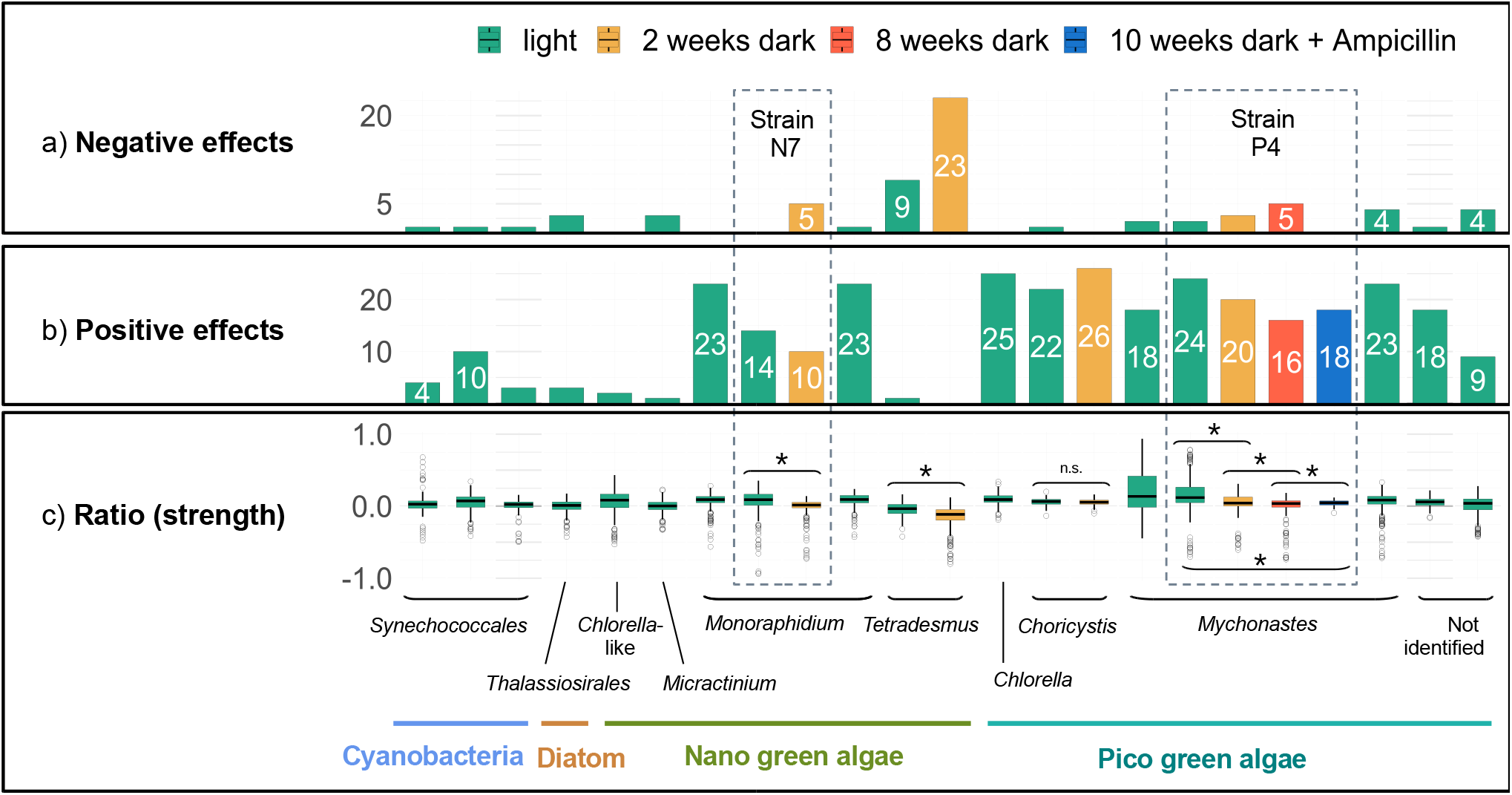
Number of significantly positive (b) and negative (a) effects (t-test, p ≤ 0.05, tab. S3) of organic compounds on the growth of phytoplankton and effect ratio (strength) (c) across three plate replicates per strain and light treatment. The x-axis shows the taxa, including various strains in some cases, and light treatment (indicated by colour). E.g. we included three different strains of *Mychonastes*, while one of them – P4 – was included with different light treatments. Strain names N7 and P4 are shown explicitly as needed for the further description. In (a) labels with less than four significant effects were removed for clarity. (c) includes all effects independent of their significance per strain and light treatment, i.e. every single measurement normalised by the control of the respective plate. This is presented by relative values around 0. Each boxplot consists of 279 values (31 substances x 3 well replicates x 3 plate replicates). Where relevant, we highlighted where the ratios of the different light treatment groups were significantly different (t-test, p ≤ 0.05) in (c).

In the same way as described, we calculated the number of significant effects and the effect strength per type of organic compound (e.g. carbohydrate, amino acids) summarised across all different strains in the standard light treatment (fig. 3a-c). We moreover indicated by ‘+’ and ‘-’ how the number of significant effects changed from light to the two weeks dark treatment with respect to the four strains included in that part of the experiment, i.e. *Mychonastes* (strain P4), *Tetradesmus, Choricystis* and *Monoraphidium* (strain N7) (fig. 3a+b).

**Fig. 3:**
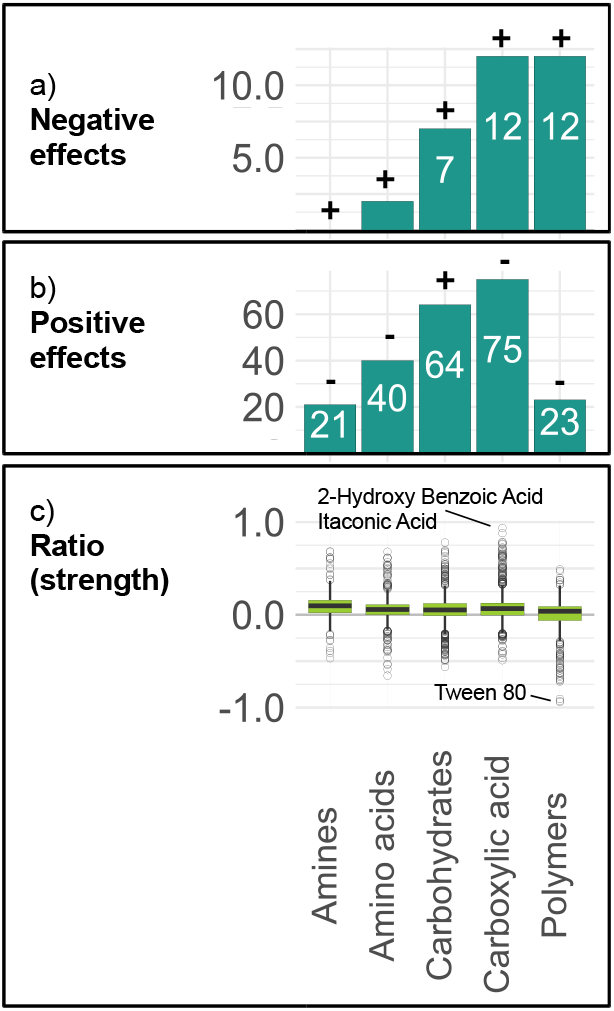
Number of significantly positive (b) and negative (a) effects (t-test, p ≤ 0.05, tab. S3) of different types of organic compounds on the growth of the 17 phytoplankton strains in the light (three plates per strain) and effect ratio (strength) (c). The ‘+’ and ‘-’ indicate how the number of significant effects changed from light to dark (two weeks incubation), referring to the four strains included in the dark experiment, namely *Mychonastes* (strain P4), *Tetradesmus, Choricystis* and *Monoraphidium* (strain N7). Precisely, ‘+’ indicates an increase of significant effects from light to dark and ‘-’ indicates a decrease of significant effects from light to dark. Note the varying numbers of compounds included per substrate type described in the methods. (c) includes all effects independent of their significance per strain and light treatment, i.e. every single measurement normalised by the control of the respective plate. This is presented by relative values around 0. The number of data points included in the box plots varied from ca. 300 to ca. 1500 depending on the number of substrates included per substrate type.

Lastly, we used the t-test to assess if the ratio was different between the different light treatments for the respective selection of strains (fig. 2c) (p ≤ 0.05, see supplementary data tab. S3).

## 2 Results

### Organic carbon drives growth in phytoplankton in the light

In the light, significantly positive effects of organic compounds appeared for all 17 strains and significantly negative effects for 13 strains (fig. 2a+b, tab. S3). While most strains experienced both positive and negative effects depending on the organic compound, the vast majority of effects were positive.

Overall, the highest number of significantly positive effects in the light treatment -up to 25 out of 31 -appeared for the pico green algae strains, particularly the three *Mychonastes* strains, *Chlorella, Choricystis* and one of the unidentified pico green algae strains (fig. 2b, tab. S3). In the group of nano green algae results varied with a low number of significantly positive effects in *Tetradesmus*, the *Chlorella*-like strain and *Micractinium* and higher numbers of 14 -23 significantly positive effects in the three *Monoraphidium* strains. The number of significantly positive effects on our diatom and the cyanobacteria were overall rather low.

Strongest positive effects in the light treatment appeared for *Mychonastes* with up to 90 % increased cell numbers, followed by the *Synechococcales* with up to 70 % increased cell numbers compared to the control (fig. 2c).

The number of significantly negative effects was overall low compared with the number of significantly positive effects under the standard light conditions (fig. 2a+b, tab. S3). The highest number of significantly negative effects here was found for *Tetradesmus* which was negatively affected by nine organic compounds.

Strongest negative effects in the light appeared for the *Monoraphidium* strain N7, as well as different *Mychonastes* strains where cell counts were diminished by up to over 90 % and up to over 70 % respectively (fig. 2c) compared to the control without organic.

### Darkness downgrades effects of organic carbon

Incubation in the dark had largely negative effects on the use of organic compounds for growth compared to the standard light conditions (fig. 2). Precisely, *Mychonastes* (strain P4), *Tetradesmus* and *Monoraphidium* (strain N7) had more significantly negative and fewer significantly positive effects after two weeks in the dark (fig. 2a+b, tab. S3). In the three strains, the overall ratio between organic treatments and controls was significantly lower after two weeks in the dark than in the light (fig. 2c, tab. S3). *Tetradesmus* was significantly negatively affected by 23 compounds in the dark (fig. 2a, tab. S3), which was the highest number of negative effects achieved by a single strain in the whole experiment. The taxon also achieved the strongest negative effects in the dark with up to 80 % reduced cell counts compared to the control (fig. 2c).

*Choricystis* achieved a higher number of significantly positive and lower number of significantly negative effects in the dark compared to the standard light conditions (fig. 2a+b, tab. S3). The overall ratio between organic treatments and the control across compounds remained identical (fig. 2c, tab. S3) indicating that there was a shift in how different substances affected the strain.

A longer dark acclimation time of eight weeks tested on *Mychonastes* (strain P4) did not positively affect the ability to use organic carbon for growth compared to the two weeks acclimation in the dark (fig. 2). Indeed, the longer acclimation time resulted in even less significantly positive and more significantly negative effects (fig. 2a+b, tab. S3) and the overall ratio between organic treatments and controls was significantly lower (fig. 2c, tab. S3).

### Ampicillin removes negative effects of organic carbon in the dark

The treatment of *Mychonastes* (strain P4) with Ampicillin after about ten weeks incubation in the dark erased all significantly negative effects on the strain (fig. 2a, tab. S3). Specifically the strong negative effects of Tween 40 and Tween 80 were removed. Due to the absence of significantly negative effects, the overall ratio between organic carbon treatments and controls was significantly higher than in the eight weeks of dark treatment without Ampicillin (fig. 2c, tab. S3). However, positive effects remained constrained compared to the light treatment (2b+c, tab. S3).

### Positive effects are caused by a high variety of different compounds

In the light, the highest number of significantly positive effects across strains was achieved by different carboxylic acids and carbohydrates, which are also the two groups with the most substrates included (fig. 3b, tab. S3). Strongest positive effects were achieved by carboxylic acids, specifically 2-Hydroxy Benzoic Acid as well as Itaconic Acid, with up to 90 % increased cell numbers (fig. 3c).

Significantly negative effects were often associated with the polymers Tween 40 and Tween 80, as well as the carboxylic acid D-Galactonic Acid g-Lactone and the carbohydrate D-Xylose (fig. 3a, tab. S3). Strongest negative effects appeared for Tween 80 where cell count was up to ca. 94 % lower than in the control (fig. 3c).

In the dark, the number of significantly negative effects overall increased for all types of compounds (fig. 3a) for the subset of strains included, i.e. *Mychonastes* (strain P4), *Tetradesmus, Monoraphidium* (strain N7) and *Choricystis*. On average it increased by about four significantly negative effects. The change was remarkably strong for carbohydrates, where ten times more significantly negative effects appeared in the dark compared to light, mostly associated with *Tetradesmus*. The number of significantly positive effects declined for all types of compounds except carbohydrates (fig. 3b), where it increased from 17 to 19. This was associated with more significantly positive effects by carbohydrates on the growth of *Mychonastes* and *Choricystis* in the dark.

## 3 Discussion

### Pico green algae and Monoraphidium particularly benefit from organic carbon

In our study, we investigated the ability of 17 phytoplankton strains to use 31 dissolved organic carbon sources for growth. All of our strains took advantage of the presence of certain organic compounds (fig. 2b). This is in compliance with former studies conducted with phytoplankton strains from the same or a related taxonomic group both from natural habitats and culture collections, including e.g. *Mychonastes* (39), *Choricystis* (40), *Monoraphidium* (41), *Tetradesmus* (42) and *Micractinium* (43). Our results support the idea that mixotrophy might be a default strategy in phytoplankton rather than an exceptional ability. Strains that were positively affected by a high number of different compounds, i.e. more than 10 to more than 20, moreover originated from all different sampling stations and seasons included (see also supplementary data tab. S1). Hence, mixotrophy seems to appear across seasons and space in the Elbe estuary.

Pico-sized green algae such as *Mychonastes, Choricystis* and *Chlorella* were significantly positively affected by a high variety of organic substances, and included some of the strongest effects (fig. 2b+c, tab. S3). Their small size allows for rapid uptake of dissolved compounds. Moreover, these fast-living creatures might be able to more quickly adjust to the availability of organic compounds than larger taxa, and show their full potential within the relatively short 24 h experiments. However, the results show that size was not necessarily the only key driver, as the larger *Monoraphidium* taxa experienced similar effects on growth as the pico greens, while the small cyanobacteria did not.

The finding of poor positive effects of organic carbon on the growth of some taxa does not necessarily imply that there were no effects at all. Other studies have shown that organic carbon can alter traits beyond growth, such as cell size (7) that were not determined in our study. Ultimately, some of the strains with a rather low number of less than 5 significantly positive effects (fig. 2b) had more significant effects on single plates which did not re-appear in the other plate replicates and were therefore not significant when the plate data were merged. Those strains seem to use more organic carbon sources under certain conditions, in this case mostly in the later exponential phase. The role of the growth phases for organic carbon acquisition has rarely been discussed to our knowledge (5). Our results highlight the necessity to integrate the life cycle into mixotrophy related research, especially taking into account that in the field taxa might often not grow exponentially.

### Organic carbon acquisition is not primarily a strategy to deal with darkness

Combining autotrophic and heterotrophic mechanisms might enable phytoplankton to maintain growth, especially under highly variable and partially stressful conditions such as light limitation, which is common in estuaries and similar habitats (20). Different studies have found mixotrophy to be associated with reduced light availability, which has largely been investigated for phagotrophs (9,23) and benthic diatoms (24). Some phytoplankton taxa have been shown to even survive complete darkness on the basis of organic carbon (4,25).

However, for the strains included in our dark experiment, we found overall less significant and weaker positive and more significant and partially stronger negative effects in the dark than in the light (fig. 2a-c, tab. S3). While significantly negative effects could be removed by addition of Ampicillin, indicating bacteria played a role here, positive effects remained constrained both in the Ampicillin treatment and with enhanced dark acclimation time.

This indicates that the poorer ability to utilise organic carbon for growth was an effect of the reduced light availability. Light dependence of organic carbon uptake and utilisation has been observed before (10,44). Though many of our strains would certainly benefit from different organic compounds in their natural habitat also at reduced light availability, our results indicate that organic carbon acquisition was not primarily a strategy to maintain growth in the dark. Factors beyond growth however, such as accumulation of storage substrates (e.g. starch and lipids) that could ensure longer term survival have not been investigated in this study and could play a role. A shift in the use of different types of compounds for growth towards a higher number of significantly positive effects of carbohydrates in the dark -specifically in *Choricystis* and *Mychonastes* -might show the effort of phytoplankton to acquire an alternative source of energy where light is limited (fig. 3b, tab. S3).

### Negative effects of organic carbon might have been promoted by bacteria

Though in an overwhelming amount of cases organic compounds had positive effects on the growth of our taxa, significantly negative effects appeared for various strains. While the positive effects appeared evenly distributed across various different compounds, a high number of negative effects was associated with the polymers Tween 40 and Tween 80 as well as the carboxylic acid D-Galactonic Acid g-Lactone. The number of significant negative effects overall increased when strains were held in darkness (fig. 2a, fig. 3a). A specifically high number of significantly negative effects appeared for *Tetradesmus* (fig. 2a). Similar results were obtained in former studies with another *Tetradesmus* strain under heterotrophic conditions (45). The results of the Ampicillin treatment on *Mychonastes* (strain P4) in the dark, where all significantly negative effects were removed, gives indications that the negative effects might be largely explained by microbial processes. Such processes might play a role in hindering phytoplankton growth in the estuary. Phytoplankton strongly decline along the estuary especially during the passage through the city of Hamburg and so far this has been led back to grazing (46,47). Potential negative effects of bacterial processes should be considered.

### Positive effects of organic do not primarily depend on the microbiome

Lastly, in the Ampicillin treatment of *Mychonastes* (strain P4) (ten weeks dark), positive effects remained similar to samples that had not received an Ampicillin treatment (eight weeks dark). Some were slightly lower, which might be an effect of the slightly longer incubation in the dark, i.e. ten weeks compared to eight weeks. Though this was only investigated with a single strain in the dark, it gives indication that the use of organic carbon for growth was largely not depending on the microbiome (e.g. substrate transformation), but a direct use by phytoplankton.

## 4 Conclusion

Organic compounds promoted the growth of phytoplankton taxa from the Elbe estuary across functional groups, season and origin. The role of phytoplankton as partial heterotrophs -e.g. in trophic upgrading -should be considered in upcoming research.

Our results moreover indicate that organic carbon acquisition is not primarily a strategy to deal with reduced light availability. However, effects beyond growth, e.g. use of organic compounds to accumulate storage substrates for longer term survival in the dark could play a role and have not been included in this study.

Bacteria might promote negative effects of organic carbon on the growth of phytoplankton. Those should be considered in further research and discussions about phytoplankton abundance in the Elbe estuary and similar habitats.

## Acknowledgements

We would like to thank Vanessa Russnak and the crew and captain of the research vessel Ludwig Prandtl (LP210308, LP210503, LP220228) for the organisation of the sampling cruises, Stefanie Schnell for the technical support in the laboratory, Luisa Listmann for implementing the organic carbon research at the IMF Hamburg as well as Inga Hense and Hans-Peter Grossart for the conceptual support.

## Funding

This project was funded by the Deutsche Forschungsgemeinschaft (DFG, German Research Foundation) as part of the project ‘Biota-mediated effects on Carbon cycling in Estuaries’ (407270017/RTG2530).

## Notes

### Competing Interest Statement

The authors have declared no competing interest.

