## Supplementary figure S1 for "Organic compounds drive growth in phytoplankton taxa from different functional groups"

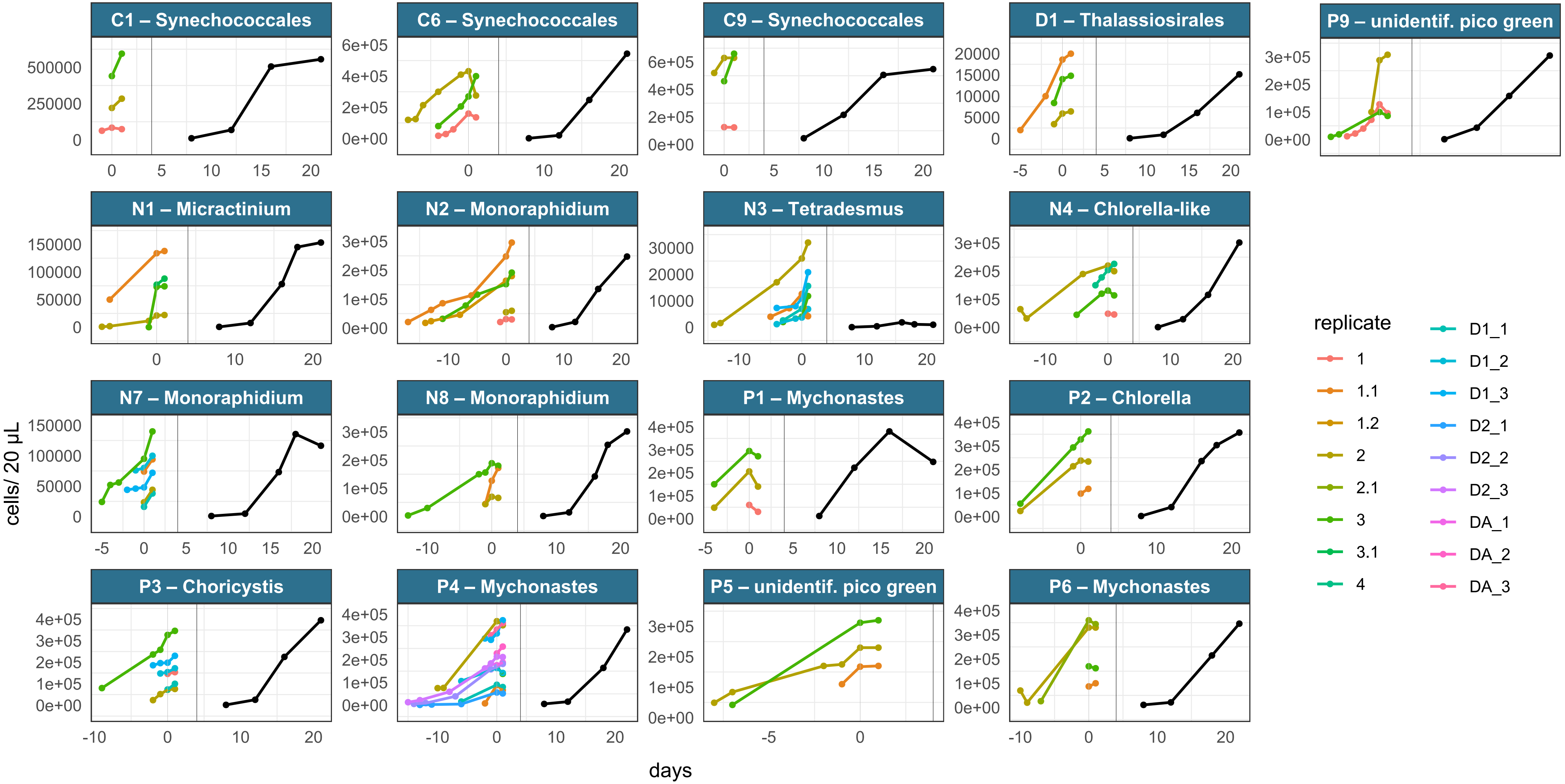


**Fig. S1: Casual tracking of cell counts of different strain cultures used for the EcoPlates™** (colours = plate replicate culture was used for; D1, D2, DA refer to the dark treatments with D1 = 2 weeks dark, D2 = 8 weeks dark, DA = 10 weeks dark + Ampicillin, see also fig. 1 and further supplementary data tab. S2. Replicates without further notice refer to standard light, some strains included more than 3 replicates in the light as shown in tab. S1 and S2). The black curve in the right panel each shows a growth curve from a former pilot studies (where available) during a 14 d period. The left side shows the cell counts from the flask cultures prior to plating (flask cultures), after 24 h in the ecoplate (day 1) and in some cases up to 2 days after plating (residue in flask). Days prior to plating are shown as respective negative values. Decline after transfer into the EcoPlates™ appears for various strains, and has been observed in a former study (Listmann et al. 2021).
